# Microbial diversity of Atlantic Rainforest ponds assessed by nanopore sequencing

**DOI:** 10.1101/2025.06.17.660208

**Authors:** Samir V. F. Atum, Douglas M. M. Soares, Isaias Santos, Grant A. Johnson, Adão H. R. Domingos, Etelvino J. H. Bechara, João C. Setubal, Cassius V. Stevani, Renato S. Freire

## Abstract

Despite their limited size, ponds are ecologically important habitats that harbor rich microbial communities and play key roles in global biodiversity and biogeochemical cycling. In this study, we conducted a metagenomic survey of microbial diversity in three ponds - Vermelha, Grande, and Furnas - situated within the Atlantic Rainforest biome in Brazil. Vermelha and Grande are natural, remote, and minimally impacted by human activity, whereas Furnas is an artificial pond with greater human accessibility. Using a long-read nanopore shotgun metagenomics approach, we sequenced DNA extracted from pond water, assembled metagenomes, and performed taxonomic classification and functional annotation. Furthermore, 21 metagenome-assembled genomes (MAGs) were recovered. Our results reveal striking biodiversity contained within ponds, with pond-specific microbial community structures and functional profiles. In addition to bacterial communities, some dsDNA bacteriophages and eukaryotic viruses were also detected. Functional annotation identified putative antimicrobial resistance genes (ARGs), most of them in Furnas, potentially reflecting human impact. Cyanotoxin biosynthesis genes were detected as well, predominantly in Vermelha. Our findings underscore the ecological distinctiveness of each pond and demonstrate the utility of nanopore-based metagenomics for investigating microbial biodiversity in understudied freshwater ecosystems, with implications for conservation and biotechnological exploration.

**GRAPHICAL ABSTRACT:** Through nanopore metagenomic sequencing, this work uncovered the microbiome of the 3 analysed freshwater ponds of the Brazilian Atlantic Forest. Results include the taxonomic profile present in each sample, countless genes identified and 21 MAGs recovered.

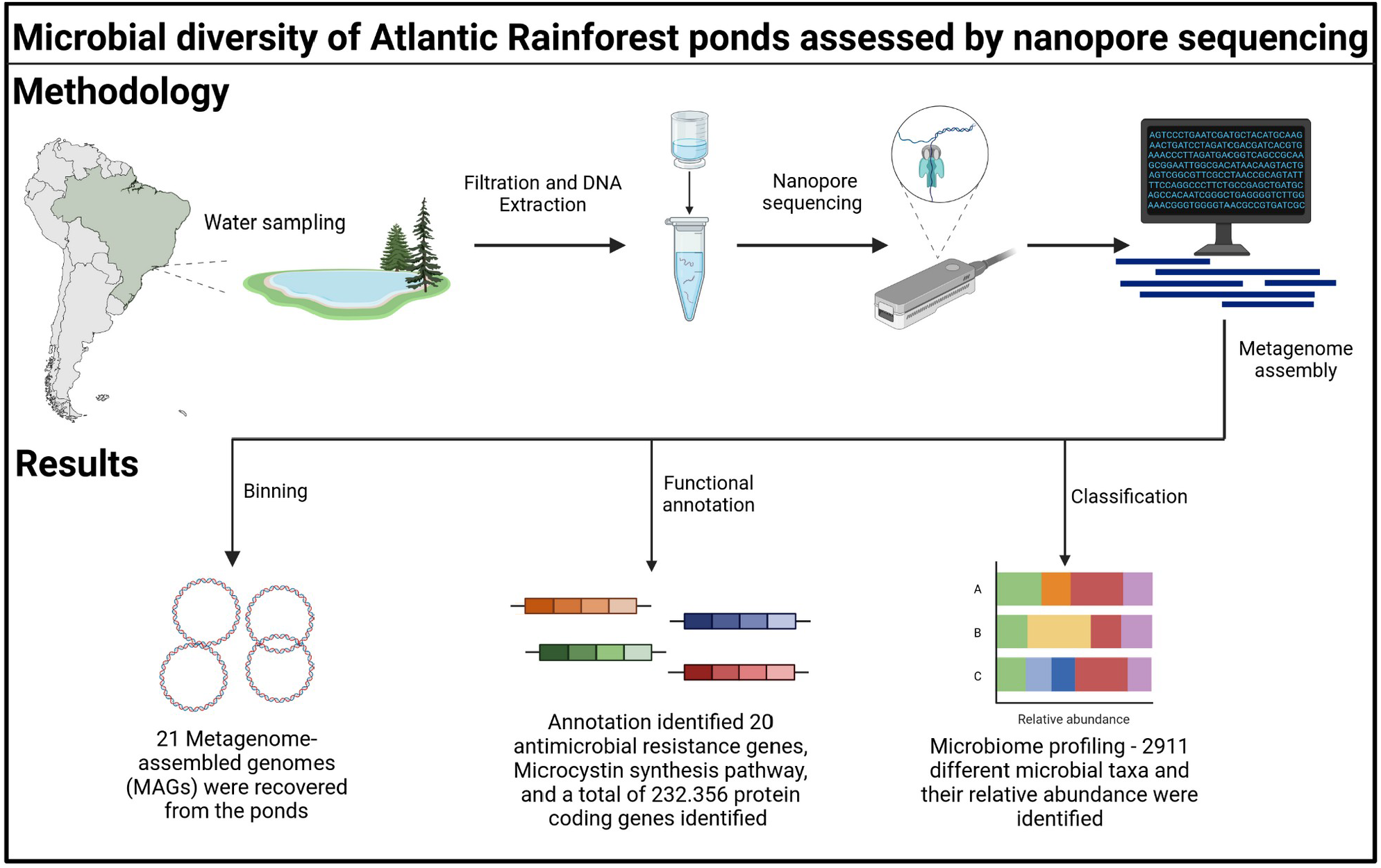

## 1. Introduction

Ponds hold significant value as microorganism reservoirs, fulfilling crucial roles in ecosystems and human activities. Despite their typical size of less than 50,000 m^2^; collectively, these water bodies globally surpass lakes in total area (Chopyk et al., 2020; Downing, 2010). With distinct ecological importance, ponds play a substantial role in the global carbon cycle and overall biodiversity (Biggs et al., 2017). They nurture diverse macro- and microspecies, while also fostering intricate microbial communities crucial for aquatic ecosystem health. However, their significance is often overshadowed by larger aquatic systems, despite being more sensitive to environmental and human influences due to their shallow nature and limited size. This sensitivity can lead to issues such as algal blooms and contamination, impacting water quality and ecosystem health (Chopyk et al., 2020). Therefore, comprehending and addressing pond microbial dynamics is vital to maintain their ecological integrity and services. Additionally, human-induced alterations in landscape dynamics, including urbanization, agricultural practices, and hydrological changes, pose threats to pond ecosystems, including nutrient loading, contamination, habitat loss, and species encroachment, with potential cascading effects on entire ecosystems, and negatively impacting plant and animal diversity, including unique and endangered species like amphibians, dragonflies, and aquatic plants that rely on ponds as refuges (Oertli et al., 2005). Studying pond microbial biodiversity can inform conservation and restoration efforts, benefiting both microbial communities and dependent species. Hence, mitigating human-induced impacts on pond ecosystems is crucial for preserving ecological function and the diversity of inhabitants.

Among the currently available tools to assess biodiversity, metagenomic approaches consist of a reliable and affordable way to identify species at a molecular level. Many metagenomic studies are based on the sequencing of DNA barcodes, such as the 16S rRNA gene and the Internal Transcribed Spacer (ITS), widely used for molecular taxonomy. While this approach is useful for exploring the diversity and microbial dynamics in environmental samples, it often falls short in revealing the potential genetic resources of the organisms and achieving high taxonomic resolution, especially when only sequencing a few variable regions of these marker genes. Additionally, its effectiveness is dependent on the completeness of databases, particularly when not using de novo assemblies.

While short-read platforms such as Illumina are renowned for high accuracy, coverage, and data throughput, they require a high initial cost, and short-reads result in highly fragmented assemblies. In this scenario, nanopore long-read sequencers are an affordable alternative to produce more contiguous assemblies. The long read technology also overcomes the uneven coverage of reads as a result of GC bias, which can be introduced during the amplification and sequencing steps of Illumina sequencing (Chen et al., 2013). Consequently, the better reads classification results in a powerful tool to assess the relative abundance of taxa (Meslier et al., 2022). Considering the necessity of *de novo* assemblies for unexplored water resources, long-read sequencing approaches offer a reliable, affordable, and informative way to investigate the microbial diversity of the ponds.

The Upper Ribeira Tourist State Park (Parque Estadual Turístico do Alto Ribeira - PETAR) is a conservation area located in the southern part of São Paulo state, and it plays a crucial role for the preservation of the Atlantic Rainforest biome. Grande pond is localized in the state park boundaries and is accessible only by 2-h hiking from Bairro da Serra (Iporanga, SP) through dense vegetation. In the case of Vermelha pond (Apiaí, SP) it is located on a wilderness plateau accessible by helicopter or a 10-h challenging hike (Fig. 1). These difficulties to reach them limit their access and utilization by humans. As a result, they remain less affected by direct human impact stemming from productive or recreational activities. Although the ponds are remote, they have already been the subject of a geological study by (Saia, 2006), who landed in Vermelha with a helicopter and analyzed the sediments of both ponds, estimating their ages at approximately 4,500 and 1,000 years, respectively. Additionally, the authors proposed that the area has been covered by vegetation for at least *ca*. 16,000 years, and the climate has remained stable and humid for the past 4,500 years (Saia, 2006). These ponds have therefore been evolving naturally for thousands of years with minimal human intervention, making them ideal subjects for microbiomics. In contrast, Furnas pond was artificially built by a mining company probably in the 1950s and is easily accessible.

**Figure 1.**
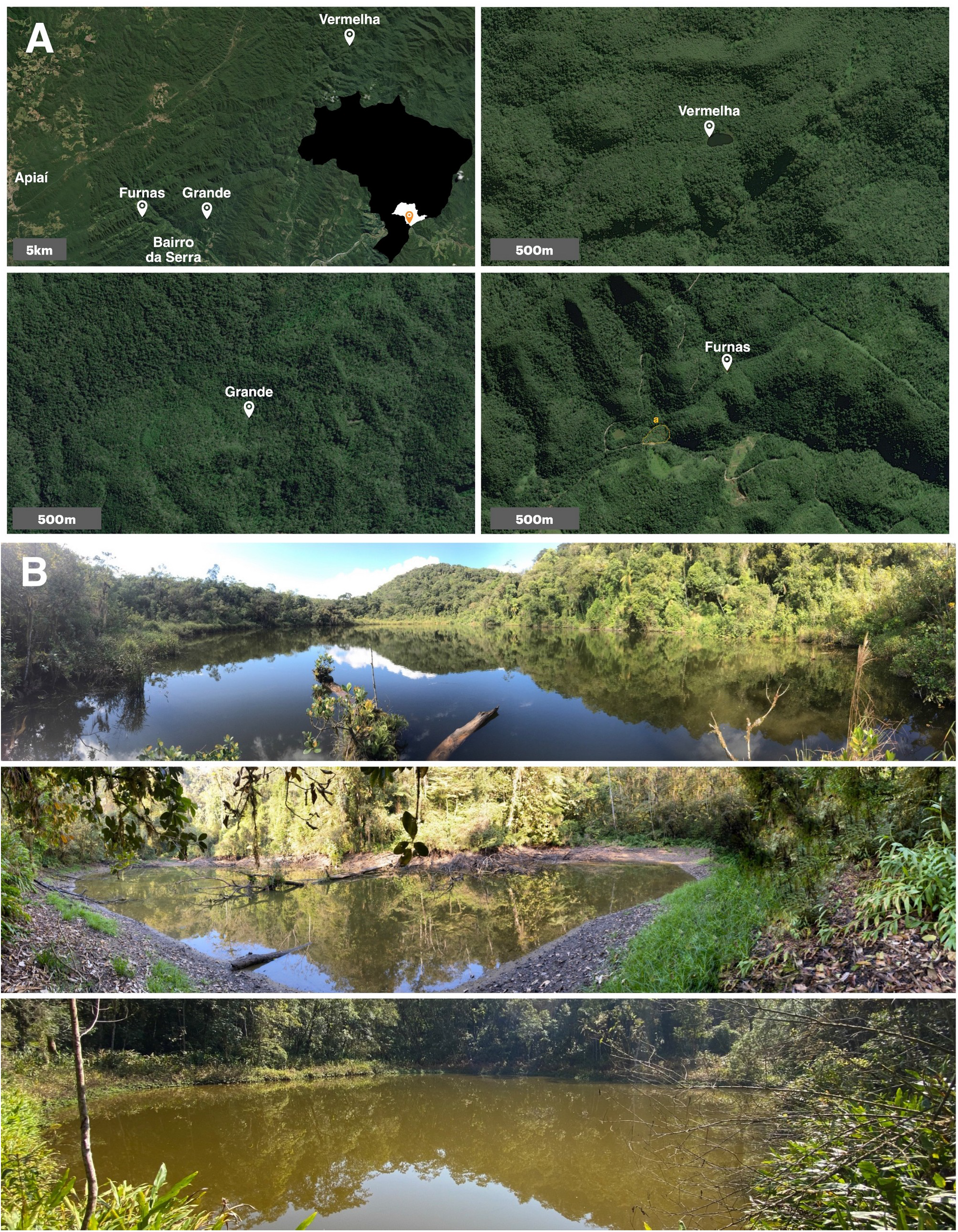
(A) Location and distance of the ponds (Vermelha, Grande and Furnas) to Bairro da Serra, Iporanga, SP, Brazil (inset). Note that Furnas Pond is located around 200m of a dirty road. In the past it supplied water for worker’s houses (A). (B) Site photos of Vermelha, Grande and Furnas ponds, respectively.

Beyond enhancing our understanding of biodiversity in these aquatic ecosystems, the data generated in this study offer valuable insights for identifying potential bioindicators of ecosystem health and for uncovering novel genes encoding enzymes with promising biotechnological applications—such as those involved in bioplastic production and bioremediation. By illuminating the unique microbial profiles of ponds within the Atlantic Rainforest, this work lays the groundwork for future studies on the microbiomes of this biome, which are essential for assessing the impacts of human activity and for prospecting genes, organisms, and biochemical pathways of interest. Ultimately, these findings can inform the development of sustainable technologies and evidence-based conservation strategies for this rich and threatened ecosystem.

## 2. Materials and methods

### 2.1. Sample collection

The estimated dimensions of Vermelha Pond (VER) are 70 × 100 m (Saia, 2006), with sampling conducted at coordinates 24º23’19”S, 48º31’50”W on May 10, 2021. Furnas Pond (FUR), measuring approximately 50 × 57 m, and Grande Pond (GDE), the smallest at 10 × 30 m, were both sampled on August 18, 2021, at coordinates 24º32’6”S, 48º39’43”W and 24º32’4”S, 48º39’44”W, respectively. The region encompassing PETAR receives an average annual rainfall of approximately 1,800 mm and has a subtropical climate, characterized by warm, humid summers and cooler, drier winters. The rainy season, spanning from October to April, accounts for roughly 80% of the total annual precipitation, with average temperatures ranging from 15°C to 30°C. For each pond, 1 L of water was collected from five different points within the euphotic zone at a depth of 0.5 m. Samples were stored at 4°C and filtered within 24 hours of collection

### 2.2. Sample filtering, DNA isolation and nanopore sequencing

Water samples were vacuum filtered using membranes with 0.22 µm pore size. Following filtration, each membrane was carefully removed with sterile tweezers and transferred into a PowerWater DNA Bead Tube (QIAGEN). Total genomic DNA was extracted using the DNeasy PowerWater Kit (QIAGEN), with an additional purification step performed using DNeasy columns (QIAGEN) to eliminate potential contaminants. DNA concentration and purity were assessed with a NanoPhotometer N80 (IMPLEN) and quantified using a Qubit fluorometer (Life Technologies). Sequencing libraries were prepared from 450 ng of genomic DNA using the SQK-RAD004 Rapid Sequencing Kit (Oxford Nanopore Technologies, ONT). Sequencing was carried out on a MinION device (ONT) equipped with R10.3 FLO-MIN111 flow cells. Sequencing progress and performance were monitored using the MinKNOW software.

### 2.3. Bioinformatic analysis

Basecalling was performed using Guppy v.5.0.7 with the high-accuracy configuration model r10.3_450_bps_hac. Metagenomic assemblies were generated using metaFlye v.2.9 (Kolmogorov et al., 2019). To remove potential contaminants, contigs were screened against the NCBI UniVec database. Taxonomic classification was carried out with Kraken2 v.2.1.1 (Lu et al., 2022; Wood et al., 2019), using the “PlusPF” database, comprising of RefSeq sequences for the human genome, prokaryotes, viruses, protozoa, and fungi - downloaded on January 7, 2024. Taxon abundance was estimated by mapping reads back to the classified contigs and counting the number of reads assigned to each taxon. The script used for this step is available at: https://github.com/Sam-Tuna/CCCC. Functional gene annotation was performed using the JGI IMG/M pipeline (Chen et al., 2023), and the resulting annotated metagenomes, along with associated metadata, have been deposited in the JGI GOLD database (Mukherjee et al., 2023). Screening for possible antimicrobial resistance genes was carried out by DeepARG (Arango-Argoty et al., 2018). Binning of assembled contigs was conducted using metaBAT2 (Kang et al., 2019), CONCOCT (Alneberg et al., 2013), and MaxBin2 (Wu et al., 2016) via the metaWRAP pipeline v.1.3 (Uritskiy et al., 2018), specifically the BINNING, BIN_REFINEMENT, and REASSEMBLE_BINS modules. The pipeline was adapted for long-read data: Minimap2 v.2.17 (Li, 2018) replaced bwa_mem (Li, 2013) for read alignment, and Flye v.2.9 was used instead of SPAdes (Bankevich et al., 2012) for bin reassembly. MAG completeness and contamination were evaluated via CheckM v.1.2 (Parks et al., 2015), and good quality MAGs (>50% completeness,<10% contamination) were classified with GTDB-Tk v.1.7 (Chaumeil et al., 2022) using the r202 database. Samtools v.1.1 (Danecek et al., 2021) was employed for the processing of multiple alignment files.

## 3. Results and discussion

### 3.1. DNA sequencing parameters and taxonomic profiling

Sequencing and assembly metrics from the Vermelha (VER), Grande (GDE), and Furnas (FUR) ponds sequencing runs are given in Supplementary Table S1. All assemblies achieved an average coverage depth of approximately 9×, sufficient for downstream analyses. Among the three datasets, the VER sample yielded the lowest sequencing throughput but exhibited the highest read N50. Consequently, its assembly was the smallest in total length yet maintained a reasonable N50 value. In contrast, GDE produced the highest sequencing throughput and the largest assembly size, although it had the lowest read and assembly N50 values. The FUR sample provided the most balanced results: while its throughput did not surpass GDE, and its read N50 was lower than VER, the resulting assembly had the highest contig N50 among all three ponds (Table S1).

Across all ponds, the microbial communities were predominantly composed of bacteria, with Pseudomonadota (formerly Proteobacteria) as the most abundant phylum (Fig. 2). Although this study provides only temporal snapshots of the microbial diversity as it probably changes throughout the year, distinct variations among the ponds are evident. The VER sample stood out for its elevated relative abundance of Cyanobacteria and Planctomycetes compared to the other sites. Planctomycetes are often associated with Cyanobacteria, and diatoms (Kaboré et al., 2020), photosynthetic protists of the phylum Bacillariophyta which were also detected in this sample, albeit in much lower abundance. These differences may be partially explained by seasonal variation: VER was sampled at the beginning of the dry season, in May, whereas FUR and GDE were sampled later in the same season, in August, when the photoperiod was approximately 20 min shorter. A shorter photoperiod can lead to reduced photosynthetic activity across all ponds; however, FUR still exhibited a notable proportion of cyanobacteria, suggesting pond-specific responses to environmental cues.

**Figure 2.**
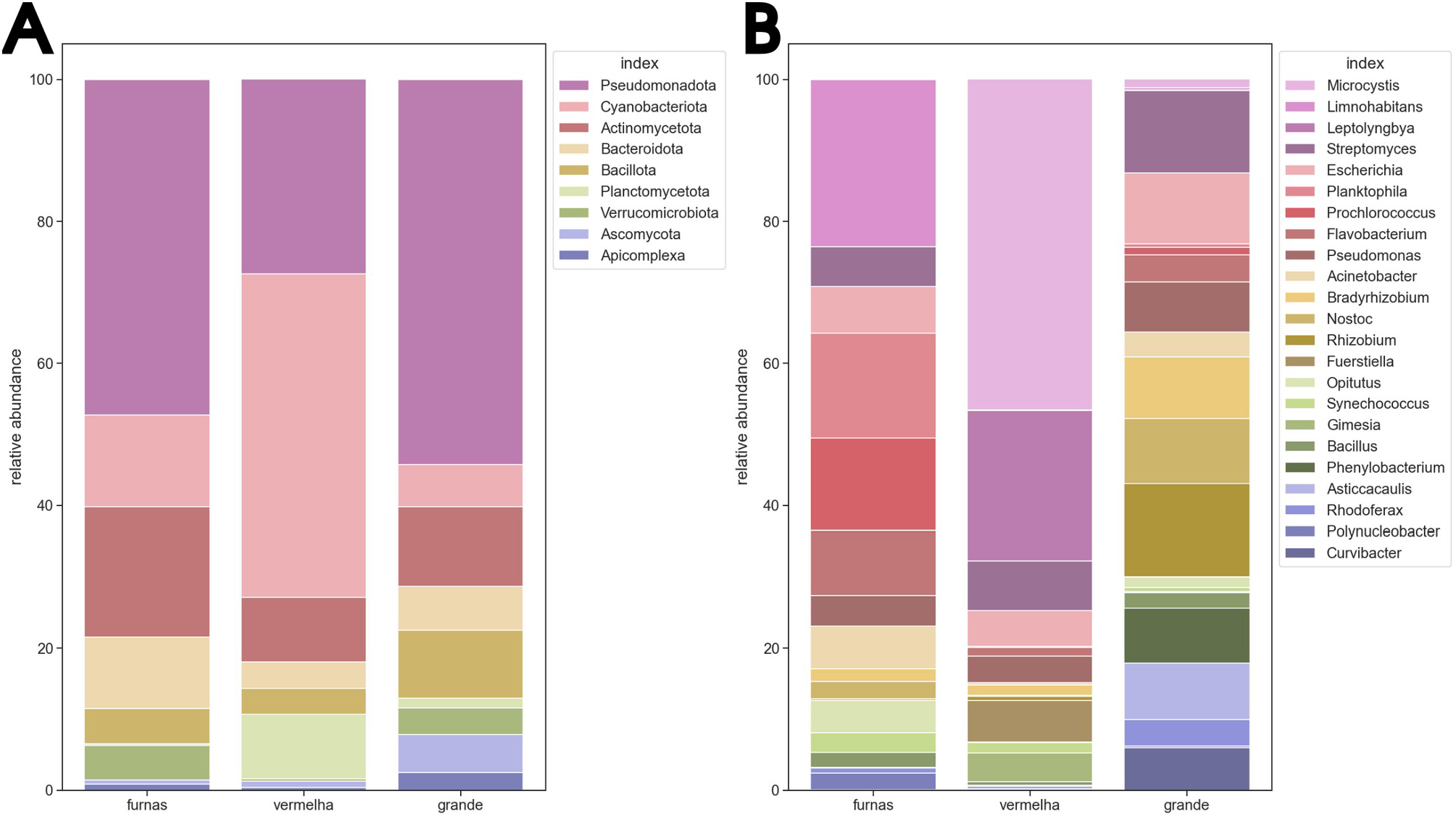
(A) Stacked bar graph showing the 10 most abundant phyla on average, across each pond. (B) Stacked bar graphs showing the 23 most abundant microbial genera. In both graphs the data was normalized and scaled within each sample so they would sum up to 100. The full phylum and genus level classification graphs are in SI (Figures S1 and S2 respectively), and the full classification results are in Supplementary File S1.

At the genus level, microbial profiles diverged further. Only three genera: *Streptomyces, Escherichia*, and *Pseudomonas* were consistently detected across all samples at comparable relative abundances. The ubiquity of these genera is notable: *Streptomyces* is renowned for its production of antimicrobial secondary metabolites (De Simeis and Serra, 2021), while *Escherichia* and *Pseudomonas* include many species with pathogenic potential. The VER pond was dominated by two genus of cyanobacteria (*Microcystis* and *Leptolyngbya*) and two Planctomycetes (*Fuerstiella* and *Gimesia*), all of which were virtually absent from the other two ponds, indicating habitat specificity. The FUR Pond displayed the highest genus-level diversity. Dominant taxa included *Limnohabitans*, a relatively understudied genus of Burkholderiales previously isolated from another pond in São Paulo state (Hahn et al., 2010); *Planktophila*, an Actinobacterium; and *Prochlorococcus*, a globally distributed cyanobacterium notably absent from VER and GDE (Biller et al., 2015). Other genera found in FUR included *Flavobacterium* (Bacteroidota), *Acinetobacter* (Gammaproteobacteria), and *Opitutus* (Verrucomicrobia), the latter represented by only one described species: *Opitutus terrae*. Finally, the microbial community in GDE pond was more evenly distributed among the most abundant genera. *Rhizobium*, whose species are generally plant root-associated, was the most abundant genus, followed by *Streptomyces, Escherichia, Nostoc* (a cyanobacterium also found in FUR but curiously absent in VER), *Asticcacaulis* (a stalked Proteobacterium), and a set of other genera including *Phenylobacterium, Bradyrhizobium, Curvibacter, Rhodoferax*, and *Novosphingobium*.

Although the primary focus of this study was not the targeted collection of viral particles, a substantial number of viral DNA sequences were recovered from the metagenomic data. Given the sequencing strategy employed, only double-stranded DNA (dsDNA) viruses were detected. Most of these viral sequences were classified as belonging to Uroviricota (primarily bacteriophages) and Nucleocytoviricota (eukaryotic viruses). Despite their relatively low representation in the total microbial communities, comprising 0.26% of relative abundance in GDE, 0.66% in VER, and 1.28% in FUR, these viral assemblages revealed notable differences in composition across the three ponds.

Viral genus-level diversity varied markedly between sites, with most genera detected being pond-specific. GDE exhibited the highest number of operational taxonomic units (OTUs) and the most even distribution among viral taxa, suggesting a more diverse and balanced viral community. In contrast, FUR was predominantly dominated by *Prasinovirus*, while VER showed a higher abundance of *Nodensvirus, Mimivirus*, and *Sokavirus*. Notably, *Prasinovirus* was the only viral genus consistently found across all three ponds, albeit with striking differences in relative abundance: it was highly prevalent in FUR and markedly less abundant in VER. As *Prasinovirus* is known to infect unicellular green algae (prasinophytes), its widespread presence serves as an indirect indicator of algal prevalence in these ecosystems.

In addition to the viral component, archaeal DNA was also detected, albeit in low abundance. Archaea accounted for approximately 0.1% of the total community in VER and FUR, and 0.5% in GDE. These archaeal sequences were primarily affiliated with the phylum Euryarchaeota, as well as members of the TACK superphylum, particularly Thermoproteota and Nitrososphaerota, indicating a minor but taxonomically diverse archaeal presence within these pond microbiomes.

### 3.2. Metagenome-Assembled Genomes (MAGs)

Although most metagenomic reads were taxonomically assigned to Pseudomonadota, the MAGs encompassed a broader phylogenetic spectrum, including members of Dependentiae, Cyanobacteria, Chloroflexota, Chlamydiota, Planctomycetota, Actinobacteriota, Verrucomicrobiota, Patescibacteria, Eisenbacteria, and Bacteroidota (Table 1). Among the three sites, FUR exhibited the greatest MAG diversity, with the phyla Actinobacteriota, Bacteroidota, Patescibacteria, and Eisenbacteria recovered exclusively from this pond. VER also showed unique MAGs, particularly those classified as Dependentiae, Chlamydiota, and Planctomycetota, while GDE was the least diverse, yielding only four MAGs (despite having the highest sequencing throughput): three from Pseudomonadota and one from Verrucomicrobiota. No MAGs from the same taxonomic order were found in more than one site. This absence of shared orders among MAGs highlights the ecological distinctiveness of each pond and suggests microhabitat-driven divergence of microbial populations. Moreover, the number of MAGs recovered per pond was positively correlated with metagenome assembly quality, as measured by N50 values (Figure S3). FUR, which had the highest assembly N50, yielded the greatest number of MAGs, followed by VER and then GDE. This underscores the impact of sequencing and assembly quality on genome recovery from complex microbial communities.

**Table 1.**
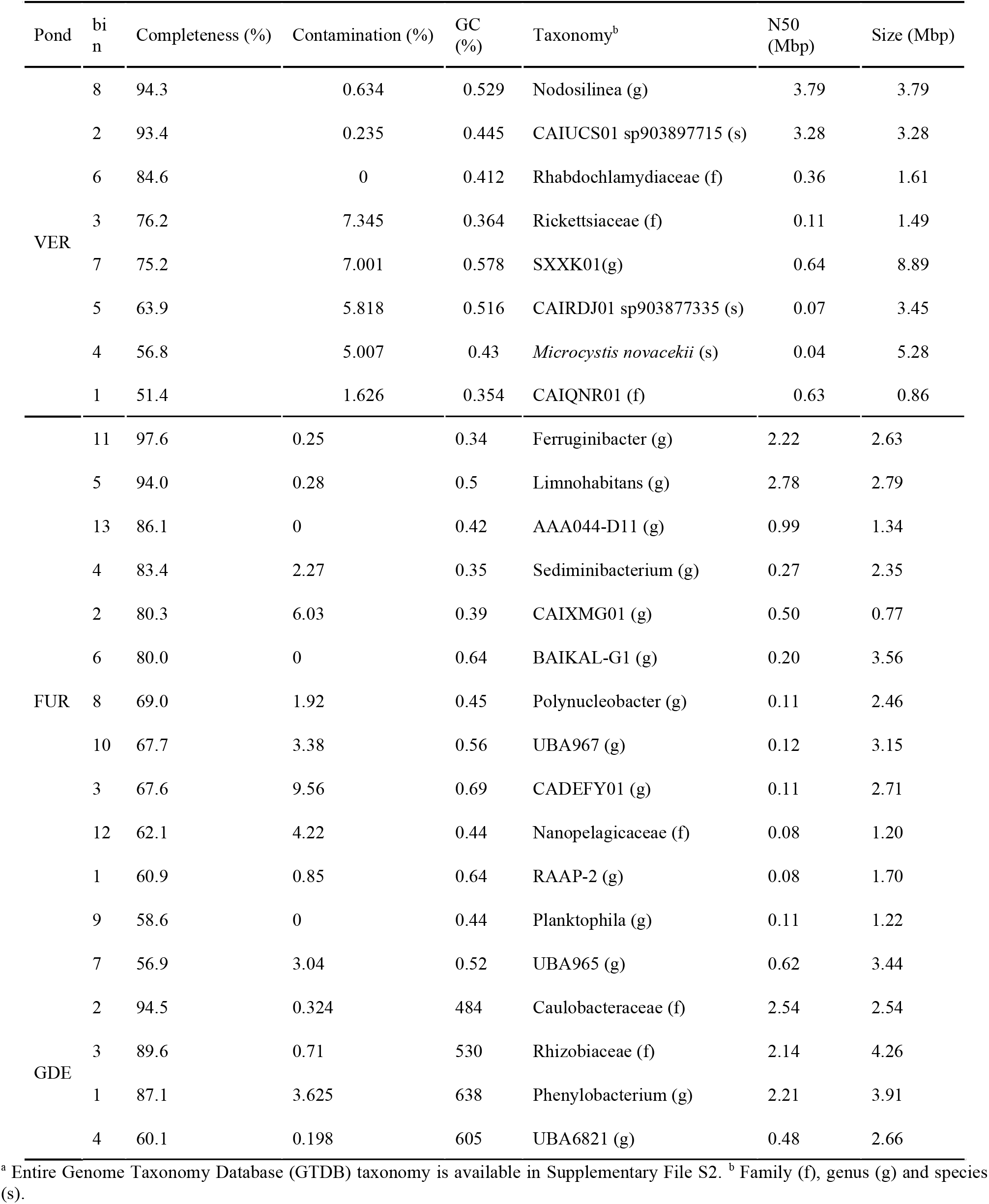
MAGs recovered from each pond.^a^.

### 3.3. Annotation

Functional annotation of the assembled metagenomes revealed a wide array of gene functions across all three ponds (File S3). Most identified genes were associated with fundamental cellular processes such as protein and nucleic acid metabolism, cell wall biosynthesis, membrane receptor signaling, and multidrug efflux mechanisms. Putative antimicrobial resistance genes (ARGs) were identified in all three ponds. FUR exhibited the highest number of ARGs, with a total of 13 genes, followed by GDE with 5, and VER with 2 (Table S2). The elevated number of resistance genes in FUR, an artificial pond with greater human accessibility, suggests a potential anthropogenic influence, possibly due to environmental contamination with antibiotic residues transported via rainwater runoff or surface flow from surrounding areas.

It is important to note, however, that antimicrobial resistance genes can also be naturally occurring, independent of human impact. For example, produced by members of the *Streptomyces* genus that was detected in all three ponds, which could be part of the reason ARGs were identified in these more remote sites. In addition to confirmed ARGs, several more general resistance-associated genes were found across all samples, including those encoding ABC-type multidrug transporters and AcrAB-TolC efflux pump components. The widespread occurrence of these resistance elements, even in relatively undisturbed environments, reinforces the importance of prudent antibiotic use and disposal practices.

Besides ARGs, the annotation also uncovered genes of environmental and biotechnological interest. These included genes involved in cyanotoxin production, bacteriocin synthesis, and polyhydroxyalkanoate (PHA) biosynthesis. In VER metagenome, the complete microcystin biosynthetic gene cluster was identified, distributed across 70 contigs. Additionally, seven contigs from the GDE assembly also contained microcystin-related genes. Regarding bacteriocins, two contigs encoding the bacteriocin *curvaticin FS47* were identified—one in VER and one in FUR. Genes involved in PHA biosynthesis were less prevalent, with only a single *phaC* gene detected, located in the VER metagenome. These findings point to the biotechnological potential harbored within these microbial communities, particularly in the context of natural product biosynthesis and sustainable biopolymer production.

Nevertheless, a substantial proportion of predicted proteins remained unclassified due to limitations in current annotation databases, which are largely biased toward either clinically significant or temperate-region organisms. Specifically, approximately 18% of predicted proteins in FUR dataset, 21% in GDE, and 23% in VER could not be functionally annotated. This underlines the ongoing challenge of studying microbial communities in underexplored environments and highlights the need to expand reference databases with genomes and genes from diverse, ecologically relevant sources.

## 4. Conclusion

The vast majority of the identified microbial DNA sequenced from the ponds is from Bacteria. While some differences between them are noticeable on a phylum level comparison, specially the heightened presence of Cyanobacteria in Vermelha, a genus level comparison better illustrates the variability between them, showcasing highly diverse microbial ecosystems.

The majority of MAGs recovered are of unknown or uncultured species, highlighting the novelty of the environments sampled, and their phylogeny was considerably more diverse than the classification results. Still, the highest abundance taxon in each pond was represented by a MAG: Vermelha pond presented a *Microcystis novacekii* genome, Furnas presented a Limnohabitans MAG and Grande had a Rhizobiaceae only classified to the family level. With the exception of the *Microcystis novacekii* genome, these are all hypothetical MAGs, since they were extracted algorithmically from the data and there are no cultured species to definitively prove their existence. However, the fact that many taxa that were not highly prevalent in the classification results were sequenced with enough coverage to generate MAGs, coupled with the fact that these MAGs could not be classified as known species, indicates that much of the microbial diversity of novel environments might go undetected even on detailed metagenomic studies, due to the databases simply not having representatives of all the sequenced taxa.

Database incompleteness is also a problem when dealing with annotation, with around 20% of proteins unclassified in each pond. Nevertheless many genes of interest were identified: all the genes involved with the biosynthetic pathway of microcystin were recovered from the Vermelha pond metagenome, coupled with one copy of the *mycC* gene from the Grande pond metagenome. A PHA synthesis gene was also present in Vermelha, and the bacteriocin curvaticin FS47 gene was identified in Vermelha and Furnas. Additionally, all three ponds exhibited a total of 20 antimicrobial resistance genes identified by deepARG, with most of them being in Furnas pond, the one with the most human contact.

It is important to note that existing genomic and functional databases are heavily biased toward microorganisms from North America and Europe. As such, the relatively remote and understudied South American ecosystems represented in this study are likely underrepresented in reference datasets. These findings underscore the importance of conducting exploratory metagenomic studies in poorly sampled biomes and reinforce the notion that, while metagenomics is a powerful tool for characterizing microbial diversity, it remains dependent on cultivated isolates and continued advances in microbial genomics to fully unlock the functional and taxonomic complexity of environmental microbiomes.

## Data availability

The raw reads obtained in this project have been deposited in the NCBI Sequence Read Archive under the accession SRR15663634, SRR33480271, and SRR33480272. The assemblies and annotations are available under the JGI GOLD project IDs Gp0614634, Gp0596633 and Gp0569555.

## Credit authorship contribution statement

SVFA: Bioinformatic analysis, graphs and statistics. DMMS: experimental design, sample preparation and sequencing. DMMS, IS, GAJ, AHRD, CVS and RSF: sample collection. SVFA, DMMS, EJHB, JCS, CVS and RSF: experimental design and draft manuscript preparation. All authors contributed to the article and approved the submitted version.

## Acknowledgments

Before peer reviewed publication, a draft of this manuscript has been submitted to BioRxiv as a preprint (https://doi.org/10.1101/2025.06.17.660208). This work was supported by Fundação de Amparo à Pesquisa do Estado de São Paulo (FAPESP, 2017/22501-2 CVS, 2022/03548-6 RSF, 2019/12605-0 DMMS, 2021/10577-0 JCS), The Brazilian National Council for Scientific and Technological Development (CNPq, 303525/2021-5 CVS, 301623/2019-8 EJHB, 305961/2019-5 JCS), the Brazilian Federal Agency for Support and Evaluation of Graduate Education (CAPES, 88887.514039/2020-00 and 88887.836021/2023-00 SVFA) and Helmholz Information & Data Science Academy (HiDA) to SVFA). We also gratefully acknowledge the support of the RCGI – Research Centre for Greenhouse Gas Innovation, hosted by the University of São Paulo (USP), sponsored by FAPESP and Shell Brasil under grants 2014/50279-4 and 2020/15230-5 (for CVS and RSF), and the strategic importance of the support given by ANP (Brazil’s National Oil, Natural Gas and Biofuels Agency) through the R&D levy regulation. The graphical abstract created in BioRender: https://BioRender.com/pv6772p. Furthermore, we thank professor Ana Lúcia Brandimarte from University of São Paulo’s Biosciences Institute for fruitful discussions and help with ecology data that sadly did not make into the paper.

