## Supplemental information for "Microbial diversity of Atlantic Rainforest ponds assessed by nanopore sequencing"

### SUPPLEMENTARY INFORMATION

#### Table S1

General data on the samples sequenced.

| | POND | | |
|---|---|---|---|
| | VER | FUR | GDE |
| Number of reads | 939,898 | 1,636,264 | 2,104,917 |
| Read N50 (bp) | 16,789 | 12,441 | 9,069 |
| Total size (bp) | 6,976,467,162 | 9,209,418,719 | 11,119,519,599 |
| Number of contigs | 7,117 | 10,157 | 28,966 |
| Contig N50 (bp) | 25,878 | 36,441 | 15,309 |
| Mean assembly coverage | 9X | 9X | 9X |
| Total assembly size (bp) | 101,980,750 | 134,224,588 | 221,908,212 |

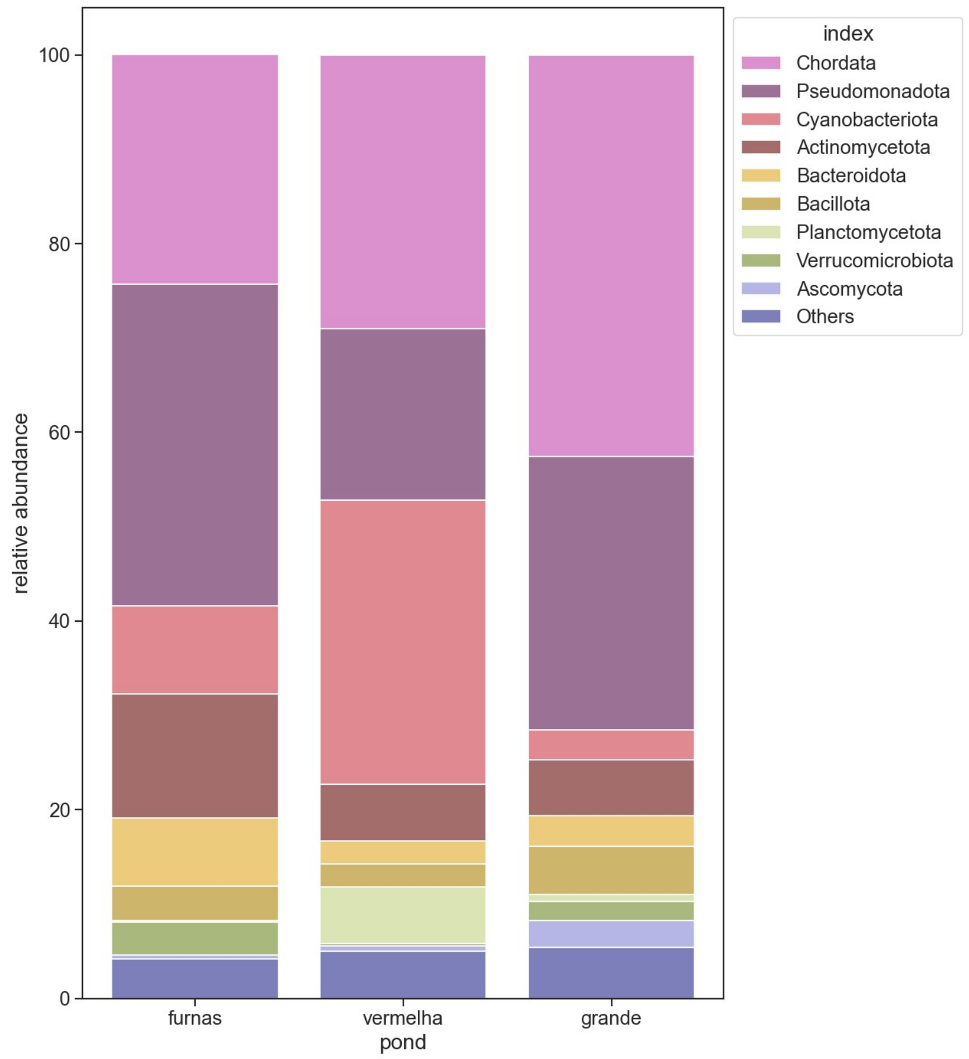


**Figure S1**: Bargraph showing the full phylum level classification of the ponds. Genera with less than 1% relative abundance are included in "Others". Full results are in Supplementary File 1.

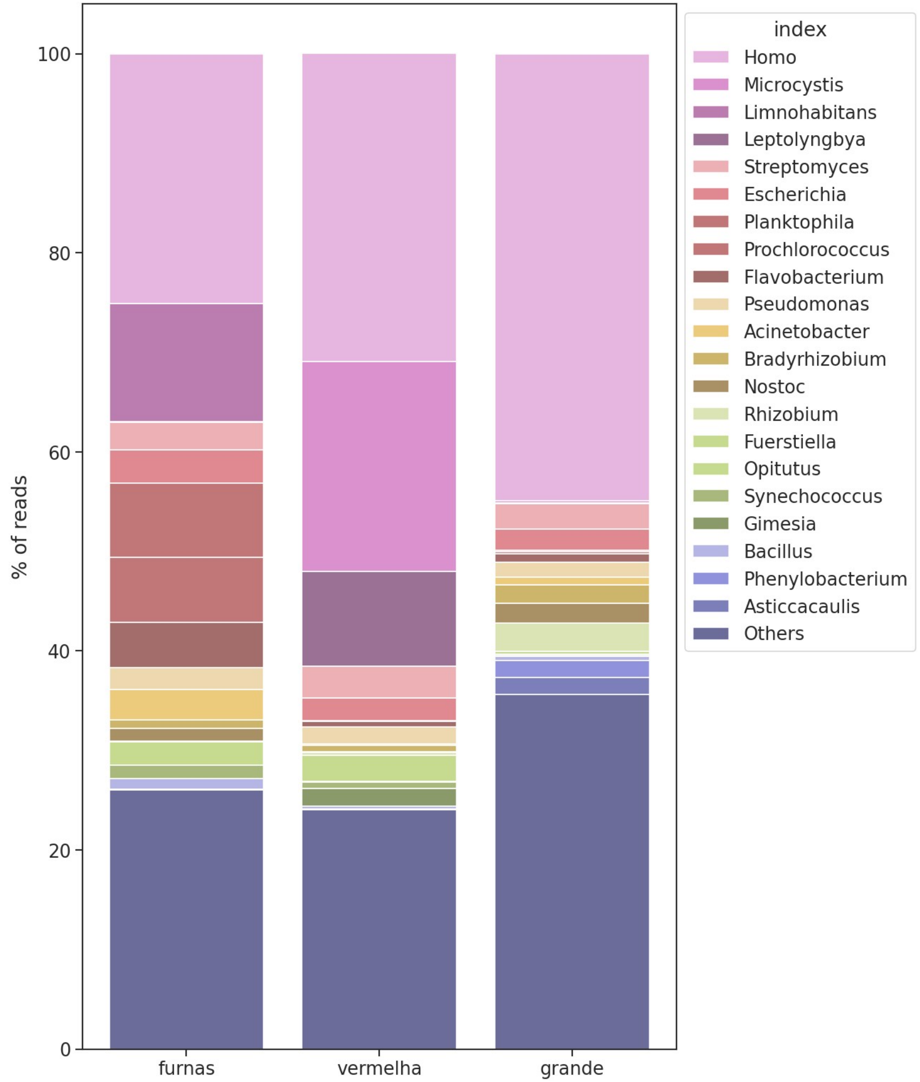


**Figure S2**: Bargraph showing the full genus level classification of the ponds. Genera with less than 1% relative abundance are included in "Others". Full results are in Supplementary File 1.

7

5
6

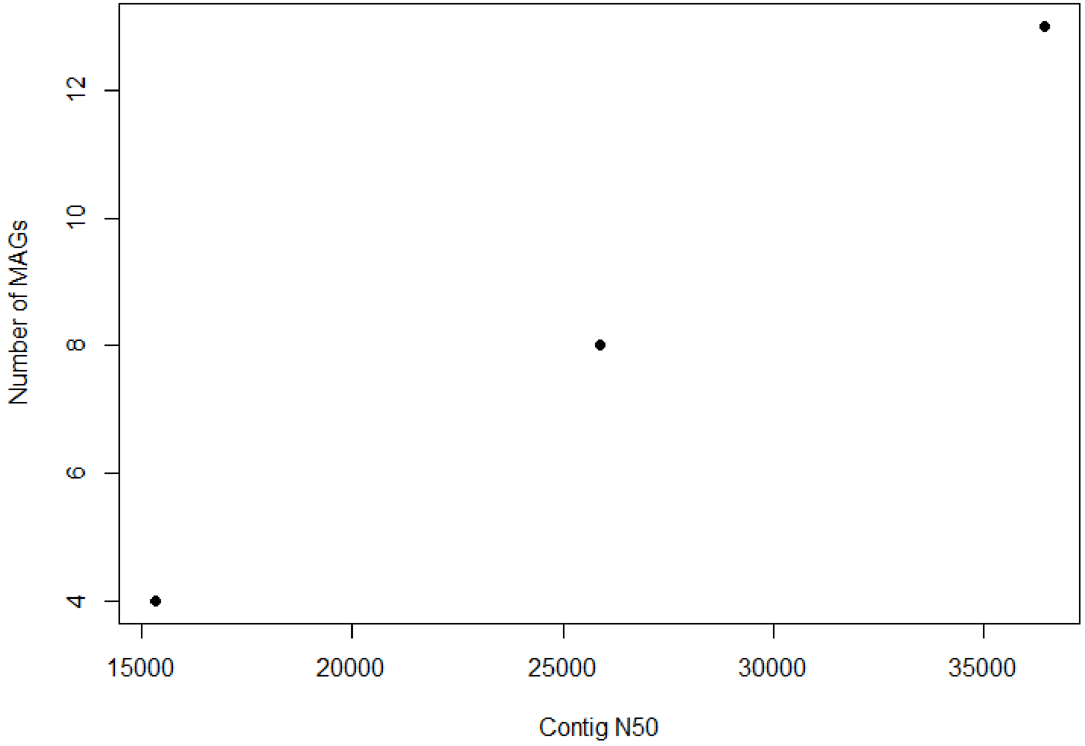


**Figure S3.** Plot of number of MAGs recovered by assembly contig N50.

**Table S2.** Putative resistance genes identified in each pond by DeepARG.

| **Pond** | **#ARG** | **contig** | **predicted_ARG-class** | **Probability** |
|---|---|---|---|---|
| VER | QACH | contig_1095 | fluoroquinolone | 0.8807206111 |
| | RPOB2 | contig_355 | multidrug | 0.9953646849 |
| FUR | RPOB2 | contig_1226 | multidrug | 0.9779076886 |
| | RPOB2 | contig_1040 | multidrug | 0.9938115669 |
| | OMPR | contig_955 | multidrug | 0.9803304806 |
| | BACA | contig_36 | bacitracin | 0.9916128332 |
| | TETA(48) | contig_3217 | tetracycline | 0.9965416081 |
| | BACA | contig_6440 | bacitracin | 0.9791516083 |
| | TETA(48) | contig_565 | tetracycline | 0.9692793709 |
| | KDPE | contig_3505 | aminoglycoside | 0.9833964448 |
| | KDPE | contig_7680 | aminoglycoside | 0.988194436 |
| | RPOB2 | contig_293 | multidrug | 0.993876158 |
| | RPOB2 | contig_744 | multidrug | 0.9940142914 |
| | TETA(48) | contig_9806 | tetracycline | 0.9971406579 |
| | BACA | contig_1319 | bacitracin | 0.993467506 |
| GDE | ADP-RIBOSYLATING_TRANSFERASE_ARR | contig_684 | rifamycin | 0.9574575721 |
| | LLMA_23S_RIBOSOMAL_RNA_METHYLTRANSFERASE | contig_27098 | MLS | 0.9997221093 |
| | AAC(3)-I | contig_20444 | aminoglycoside | 0.9999999874 |
| | RPOB2 | contig_68 | multidrug | 0.9953646849 |
| | OMPR | contig_2702 | multidrug | 0.9965310699 |
